# Evaluation of ddRADseq for reduced representation metagenome sequencing

**DOI:** 10.1101/138883

**Authors:** Michael Liu, Paul Worden, Leigh G. Monahan, Matthew Z. DeMaere, Catherine M. Burke, Steven P. Djordjevic, Ian G. Charles, Aaron E. Darling

## Abstract

**Background:** ‘Who is doing what’ is the ultimate open question in microbiome study. Shotgun metagenomics is often applied to gain knowledge of functional roles for bacteria in microbial communities, where the data can be used to predict protein encoding genes and enzymatic pathways present in the community, sometimes leading to testable hypotheses for microbial function. We describe a method and basic analysis for a metagenomic adaptation of the double digest restriction site associated DNA sequencing (ddRADseq) protocol for reduced representation metagenome profiling. This technique takes advantage of the sequence specificity of restriction endonucleases to construct an Illumina-compatible sequencing library containing DNA fragments that are between a pair of restriction sites located within close proximity. This results in a reduced sequencing library with coverage breadth that can be tuned by size selection.

**Results:** We assessed the performance of the metagenomic ddRADseq approach by applying the method to human stool samples and generating sequence data. We evaluate the extent to which ddRADseq data provides an unbiased reduced representation for microbiome profiling.

**Conclusion:** Although ddRADseq does introduce some bias in taxonomic representation, the bias is likely to be small relative to DNA extraction bias. ddRADseq appears feasible and could have value as a tool for metagenome-wide association studies.

## BACKGROUND

‘Who is doing what’ is the ultimate open question in microbiome studies. Shotgun metagenomics is often applied to gain knowledge of functional roles for bacteria in microbial communities, where the data can be used to predict protein encoding genes and enzymatic pathways present in the community, sometimes leading to testable hypotheses for microbial function. Advances in DNA sequencing technology and computing have dramatically accelerated the development of sequence-based metagenomics, which has been proposed by many scientists as a means to characterize the function of microbes in microbiomes [1, 2]. Nevertheless, studies seeking to link microbial community phenotype to genes in the metagenome, e.g. metagenome-wide association studies (MGWAS) can require large numbers of samples to be processed. As a result, MGWAS on genetically diverse samples such as mammalian faeces and soil remains difficult or intractable, due to the prohibitive cost of shotgun metagenome sequencing to adequate depth.

Here, we investigate the potential of reduced representation sequencing to be used for low-cost metagenome profiling. We describe a metagenomic adaptation of <u>d</u>ouble <u>d</u>igest restriction site <u>a</u>ssociated <u>D</u>NA <u>seq</u>uencing (ddRADseq)[3], a method for genotyping by sequencing studies of large and complex individual genomes (e.g. plants). This approach takes advantage of the sequence specificity of restriction endonucleases, where the genomic DNA is first fragmented by restriction digestion to construct a set of sequences that are flanked by the targeted restriction sites. This results in a sequencing library with complexity that is reduced roughly in proportion to the density of the restriction enzyme cut-sites, modulo any fragment size selection. The design of this approach also includes a dual-index combinatorial barcoding approach allowing samples to be multiplexed in a single sequencing run. We demonstrate the method on human fecal samples and compare the recovered taxonomic profiles to those obtained by standard metagenome shotgun sequencing on the same samples.

## METHODS

Three healthy adult stool samples were collected and the DNA extracted using an UltraClean Microbial DNA isolation kit (MO BIO Laboratories). The ddRADseq metagenome libraries were generated following the original ddRADseq protocol [3], which uses a double restriction digest followed by size selection to construct a library containing only a defined subset of the total genomic DNA, with fine-scale control of library complexity available via size selection. Combinations of commercially available restriction enzymes were computationally evaluated for cut site density and other properties in a set of reference genomes chosen to reflect a wide range of G+C content. The evaluation suggested a short list of optimal enzyme combinations, and the combination of NlaIII and HpyCH4IV (New England BioLabs) was selected for their properties of buffer compatibility, insensitivity to *dam* methylation, overhang incompatibility, and heat sensitivity. This combination of restriction enzymes was predicted to minimise bias in the representation of individual species within the metagenomic community. The protocol employed a dual index approach for sample barcoding where each adapter carries a sample barcode compatible with standard Illumina multiplexing index reads (Figure 1). Finally, a variable length region containing randomly synthesised nucleotides is included immediately downstream of the read priming site, to improve cluster calling and offer the potential to identify PCR duplicates via unique molecular identifiers [4].

**Figure 1.**
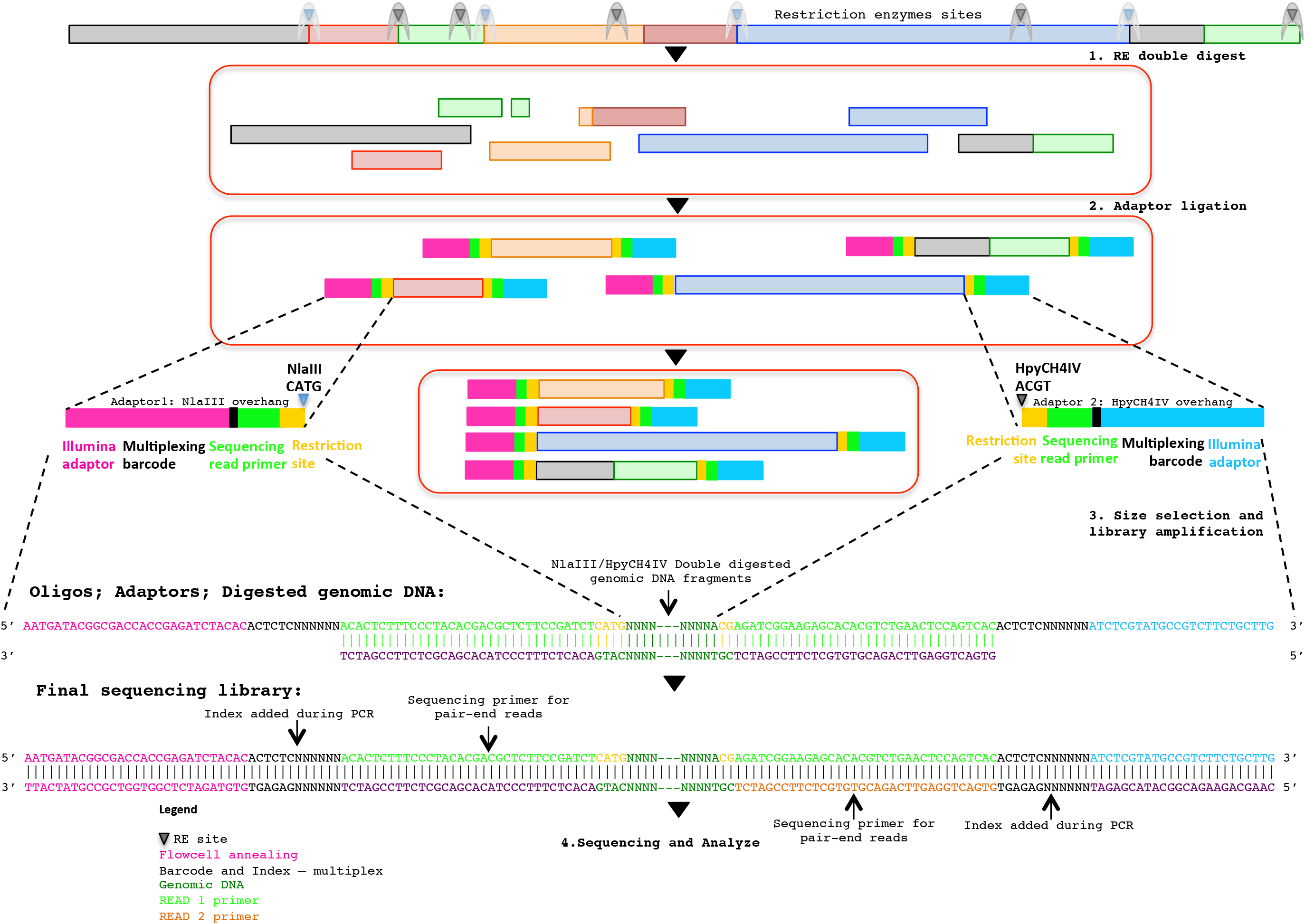
Diagrammatic representation of the ddRAD-seq method: the oligos, adaptors and final sequencing library.

To generate the ddRADseq libraries, 50ng of DNA from each sample was used in a restriction double digest following the reaction conditions recommended by the enzyme manufacturer. A total of 5U of both enzymes were used in the reaction and subsequently heat inactivated following the manufacturer's instructions. A generous molar ratio of 1:40 (digested DNA: sequencing adaptors) was then used for sequencing adaptor ligation, which ensured an excess of adaptors for ligation. Amplification of the adaptor-ligated fragments was then carried out using the Illumina standard P5 and P7 flowcell oligo primers. The three sequencing libraries were then pooled with equal volume and size selected for approximately 500-600 bp fragments with SPRI beads using a left and right side clean-up 0.5× and 0.6× respectively. Shotgun metagenomic libraries were prepared from separate aliquots of sample DNA using the Illumina Nextera DNA kit. Sequencing of those samples were done with half a lane of the Illumina HiSeq 2500 in rapid PE250 mode.

## QUALITY ASSURANCE

The quality of sequence data was examined using FastQC [5]. Reads were also checked for the presence of the sequenced portion of the restriction enzyme recognition site (NlaIII: CATG, HpyCH4IV: CGT) starting at position 0-3 in the read depending on the phasing of the barcode adapter associated with each sample.

## INITIAL FINDINGS

We present a full method for sample preparation using <u>d</u>ouble <u>d</u>igest restriction site <u>a</u>ssociated <u>D</u>NA <u>seq</u>uencing (ddRADseq) technique. Paired-end Illumina sequencing was carried out on both shotgun and ddRADseq libraries for three human gut samples. As a pilot study with only three samples, there is insufficient statistical power to evaluate associations between phenotype and microbiome genotype in these samples. Instead, we evaluate the extent to which ddRADseq yields a similar estimate of community taxonomic profile as that obtained from shotgun metagenome sequencing of the same samples. If the ddRADseq is strongly biased against (or in favour of) some taxonomic groups it might reduce (or increase) power to detect associations between phenotype and the protein coding genes in the genomes of that group. Both the shotgun and ddRADseq reads were taxonomically profiled using MetaPhlAn [6], which counts reads matching to clade-specific protein-coding marker genes in order to estimate taxon abundance from metagenomic read data. The taxonomic profile between individuals was different on both the genus and species levels, consistent with the inter-individual variation that is typically observed in human microbiome studies. Highly similar, but not identical, taxonomic profiles were observed between the shotgun-sequenced and ddRADseq metagenome libraries. Hierarchical clustering using the Manhattan distance showed that ddRADseq samples cluster closely with their matching shotgun samples (Figure 2). These results suggest that the representation of protein coding genes in ddRADseq metagenomes may be uniform enough to support association studies.

**Figure 2.**
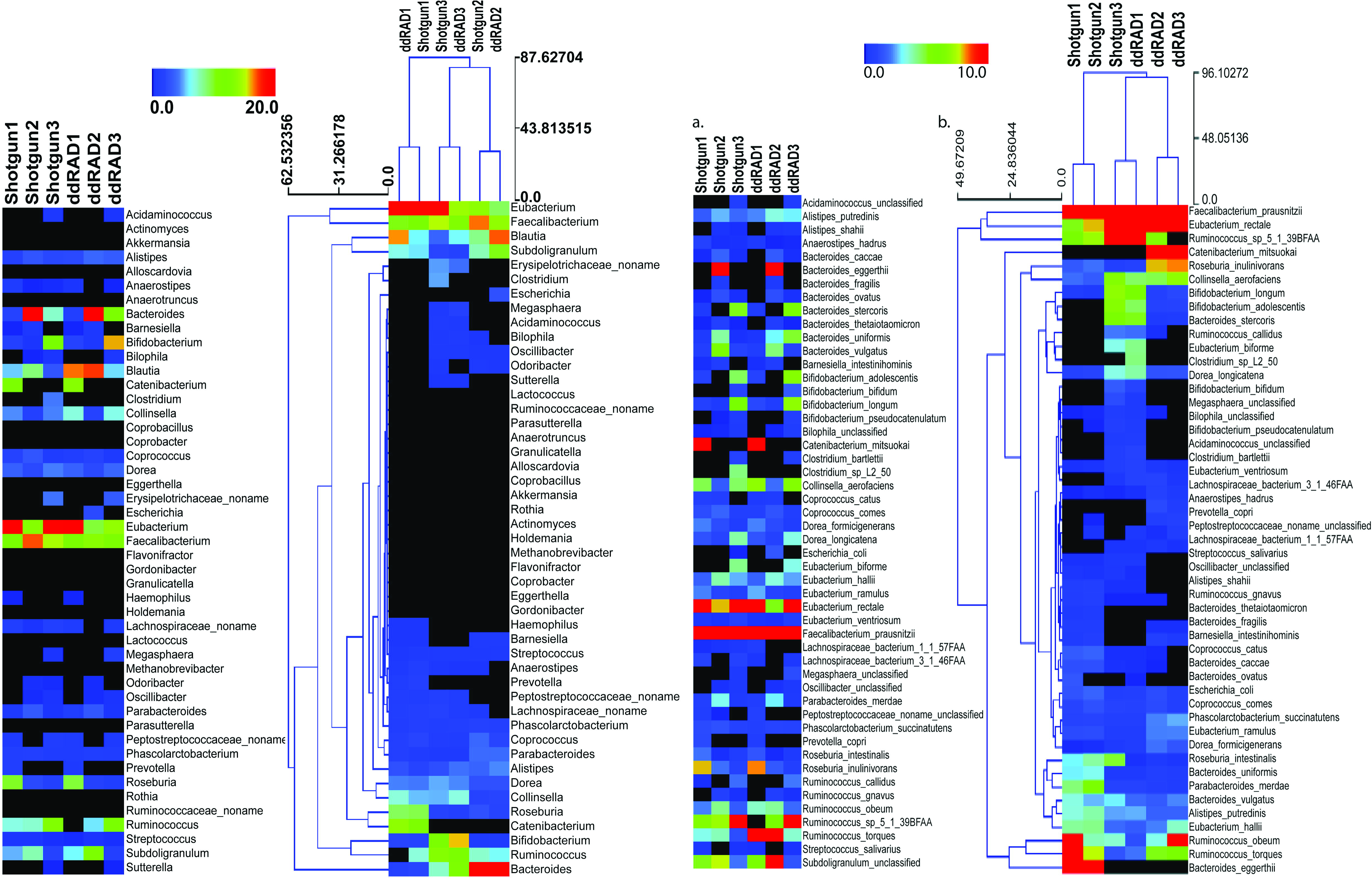
Abundance of specific phylotypes detected in all libraries on both genus (a) and species (b) levels. The heatmap is plotted and hierarchically clustered according to the abundance of each phylotype.

To assess whether the restriction enzyme pair used in this study created a reduced representation library that was biased by genomic G+C content we plotted the estimated relative abundance of each taxonomic group with respect to the genomic G+C content of a reference genome from that taxonomic group. As can be seen in Figure 3, if an association exists between genomic G+C and relative abundance in our ddRADseq libraries, it is not visually apparent. We refrain from reporting Spearman correlations on these data, as such correlation tests have well known problems on compositional data [7].

**Figure 3.**
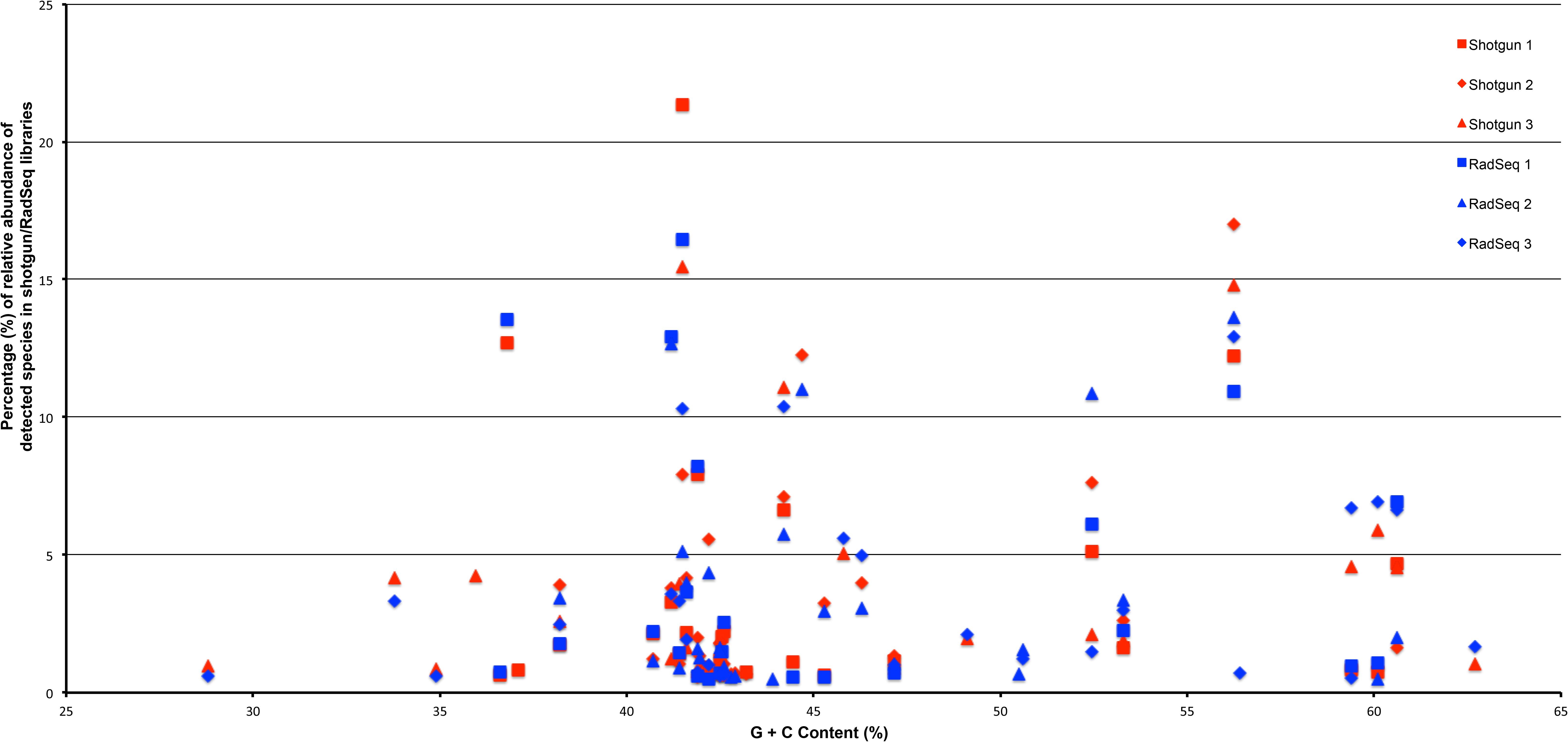
Abundance of taxonomic groups in both shotgun and ddRADseq libraries as a function of genomic GC content for all three samples. Although differences in estimated relative abundance exist between shotgun and ddRADseq libraries, the ddRADseq protocol we used has not created a bias with an obvious association to genomic GC content.

## FUTURE DIRECTIONS

Changes in the human microbiota have been associated with complex disease. The gut microbiota, for example, has been linked to conditions such as irritable bowel syndrome and inflammatory bowel disease. Major research efforts are now underway to move beyond taxonomic associations to discover and describe the genetic basis for these microbiome-associated diseases. Although shotgun metagenomics is one way to carry out those analyses, it can be prohibitively expensive in some situations. Therefore, reduced representation techniques such as ddRADseq may provide a useful means to enable large-scale studies within a fixed research budget. The reported data represents (to our knowledge) the first metagenomic adaptation of the traditional ddRADseq protocol on real microbiome samples. Here we presented a preliminary analysis of the data and showed that metagenome profiling with ddRADseq appears to be feasible. Taxon relative abundances estimated from clade-specific marker genes are similar but not identical to estimates from shotgun metagenomic data, implying that protein coding gene profiles from ddRADseq would likewise be similar to those obtained with shotgun metagenomics. When combined with careful fragment size selection, ddRADseq profiling could have value as a cost effective means to generate metagenomic profiles for use with metagenome-wide association studies and as a complementary tool for surveillance of microbial ecosystems, tracking differences across environments, treatments or time scale.

Even if enzyme choice leads to some bias in metagenomic ddRADseq libraries, other sources of bias may be more significant. For example, DNA extraction efficiency for individual community members can depend heavily on features such as the cell wall architecture of those organisms, leading to extreme bias in the extracted DNA and final sequencing library [8, 9]. The biases introduced by ddRADseq may be small in comparison to those introduced by DNA extraction. Sensitivity of restriction enzymes to methylation and other poorly understood DNA modifications maybe another factor contributing towards the observed differences in detection. The cleavage of target DNA may be blocked, or impaired, when a particular base in the enzyme's recognition site is modified. Future work to test this could explore the use of enzymes that have the same recognition site (isoschizomers), but with different methylation sensitivity, to identify modified bases through changes in the efficiency of restriction digest.

## AVAILABILITY OF SUPPORTING DATA

Shotgun sequence data are available from the NCBI Short Read Archive, project accession SRP100899, containing two datasets per sample.

### i5 NlaIII adapter oligo sequences

1. **i5 NlaIII ACTCTC** AATGATACGGCGACCACCGAGATCTACACACTCTCNNNNNNACACTCTTTCCCTACACGACGCTCTTCCGATCTCATG
2. **i5 NlaIII ATCCGG** AATGATACGGCGACCACCGAGATCTACACATCCGGNNNNNNACACTCTTTCCCTACACGACGCTCTTCCGATCTACATG
3. **i5 NlaIII GAGGAC** AATGATACGGCGACCACCGAGATCTACACGAGGACNNNNNNACACTCTTTCCCTACACGACGCTCTTCCGATCTGTCATG
4. **i5 NlaIII ACCGGC** AATGATACGGCGACCACCGAGATCTACACACCGGCNNNNNNACACTCTTTCCCTACACGACGCTCTTCCGATCTTGGCATG
5. **i5 NlaIII TGCCGT** AATGATACGGCGACCACCGAGATCTACACTGCCGTNNNNNNACACTCTTTCCCTACACGACGCTCTTCCGATCTCATG
6. **i5 NlaIII AGGCTT** AATGATACGGCGACCACCGAGATCTACACAGGCTTNNNNNNACACTCTTTCCCTACACGACGCTCTTCCGATCTACATG
7. **i5 NlaIII ATAACC** AATGATACGGCGACCACCGAGATCTACACATAACCNNNNNNACACTCTTTCCCTACACGACGCTCTTCCGATCTGTCATG
8. **i5 NlaIII GGAGGC** AATGATACGGCGACCACCGAGATCTACACGGAGGCNNNNNNACACTCTTTCCCTACACGACGCTCTTCCGATCTTGGCATG

### i5 NlaIII stub oligo sequences

1. **i5 NlaIII Phaser0 stub** AGATCGGAAGAGCGTCGTGTAGGGAAAGAGTGT
2. **i5 NlaIII Phaser1 stub** TAGATCGGAAGAGCGTCGTGTAGGGAAAGAGTGT
3. **i5 NlaIII Phaser2 stub** ACAGATCGGAAGAGCGTCGTGTAGGGAAAGAGTGT
4. **i5 NlaIII Phaser3 stub** CCAAGATCGGAAGAGCGTCGTGTAGGGAAAGAGTGT

### i7 HpyCH4IV adapter oligo sequences

1. **i7 HpyCH ACTCTC** CGAGATCGGAAGAGCACACGTCTGAACTCCAGTCACACTCTCNNNNNNATCTCGTATGCCGTCTTCTGCTTG
2. **i7 HpyCH ATCCGG** CGAAGATCGGAAGAGCACACGTCTGAACTCCAGTCACATCCGGNNNNNNATCTCGTATGCCGTCTTCTGCTTG
3. **i7 HpyCH GAGGAC** CGAGATCGGAAGAGCACACGTCTGAACTCCAGTCACGAGGACNNNNNNATCTCGTATGCCGTCTTCTGCTTG
4. **i7 HpyCH ACCGGC** CGAAGATCGGAAGAGCACACGTCTGAACTCCAGTCACACCGGCNNNNNNATCTCGTATGCCGTCTTCTGCTTG
5. **i7 HpyCH TGCCGT** CGAGATCGGAAGAGCACACGTCTGAACTCCAGTCACTGCCGTNNNNNNATCTCGTATGCCGTCTTCTGCTTG
6. **i7 HpyCH AGGCTT** CGAAGATCGGAAGAGCACACGTCTGAACTCCAGTCACAGGCTTNNNNNNATCTCGTATGCCGTCTTCTGCTTG
7. **i7 HpyCH ATAACC** CGAGATCGGAAGAGCACACGTCTGAACTCCAGTCACATAACCNNNNNNATCTCGTATGCCGTCTTCTGCTTG
8. **i7 HpyCH GGAGGC** CGAAGATCGGAAGAGCACACGTCTGAACTCCAGTCACGGAGGCNNNNNNATCTCGTATGCCGTCTTCTGCTTG
9. **i7 HpyCH CGACCT** CGAGATCGGAAGAGCACACGTCTGAACTCCAGTCACCGACCTNNNNNNATCTCGTATGCCGTCTTCTGCTTG
10. **i7 HpyCH GTCGTC** CGAAGATCGGAAGAGCACACGTCTGAACTCCAGTCACGTCGTCNNNNNNATCTCGTATGCCGTCTTCTGCTTG
11. **i7 HpyCH GATCAA** CGAGATCGGAAGAGCACACGTCTGAACTCCAGTCACGATCAANNNNNNATCTCGTATGCCGTCTTCTGCTTG
12. **i7 HpyCH GTTGCG** CGAAGATCGGAAGAGCACACGTCTGAACTCCAGTCACGTTGCGNNNNNNATCTCGTATGCCGTCTTCTGCTTG

### i7 HpyCH4IV stub oligo sequences

1. **i7 HpyCH Phaser0 stub:** GTGACTGGAGTTCAGACGTGTGCTCTTCCGATCT
2. **i7 HpyCH Phaser1 stub:** GTGACTGGAGTTCAGACGTGTGCTCTTCCGATCTT

### RADseq library amplification oligo sequences

1. **P5 primer oligo:** AATGATACGGCGACCACCGA
2. **P7 primer oligo:** CAAGCAGAAGACGGCATACGA

## DECLARATIONS

### Acknowledgements

Ethical approval for this study was obtained from the University of Technology Sydney Human Research Ethics Committee (Approval number: UTS HREC REF NO. 2014000448). We acknowledge the Centre for Digestive Disease in Sydney, NSW, Australia for their help during sample collection and for helpful discussions and advice.

### Authors’ contributions

ML, PW, LM, and CB processed the samples in the lab and did the sequencing using the Illumina MiSeq desktop sequencer. ML and AD performed the bioinformatic processing of sequence data. MZD did data analysis to predict good restriction enzyme combinations. ML and AD did the analysis, generated preliminary results, and AD uploaded the sequences to. AD, SD, and IC supervised the work. ML and AD wrote the manuscript. IC and AD designed the experiment. All authors participated in the discussions about data interpretation and manuscript supervision. All authors read and approved the final manuscript.

